# Senolytic therapy preserves blood-brain barrier integrity and promotes microglia homeostasis in a tauopathy model

**DOI:** 10.1101/2024.03.25.586662

**Authors:** Minmin Yao, Zhiliang Wei, Jonathan Scharff Nielsen, Aaron Kakazu, Yuxiao Ouyang, Ruoxuan Li, Tiffany Chu, Susanna Scafidi, Hanzhang Lu, Manisha Aggarwal, Wenzhen Duan

## Abstract

Cellular senescence, characterized by expressing the cell cycle inhibitory proteins, is evident in driving age-related diseases. Senescent cells play a crucial role in the initiation and progression of tau-mediated pathology, suggesting that targeting cell senescence offers a therapeutic potential for treating tauopathy associated diseases. This study focuses on identifying non-invasive biomarkers and validating their responses to a well-characterized senolytic therapy combining dasatinib and quercetin (D+Q), in a widely used tauopathy mouse model, PS19. We employed human-translatable MRI measures, including water extraction with phase-contrast arterial spin tagging (WEPCAST) MRI, T2 relaxation under spin tagging (TRUST), longitudinally assessed brain physiology and high-resolution structural MRI evaluated the brain regional volumes in PS19 mice. Our data reveal increased BBB permeability, decreased oxygen extraction fraction, and brain atrophy in 9-month-old PS19 mice compared to their littermate controls. (D+Q) treatment effectively preserves BBB integrity, rescues cerebral oxygen hypometabolism, attenuates brain atrophy, and alleviates tau hyperphosphorylation in PS19 mice. Mechanistically, D+Q treatment induces a shift of microglia from a disease-associated to a homeostatic state, reducing a senescence-like microglial phenotype marked by increased p16/INK4a. D+Q-treated PS19 mice exhibit enhanced cue-associated cognitive performance in the tracing fear conditioning test compared to the vehicle-treated littermates, implying improved cognitive function by D+Q treatment. Our results pave the way for application of senolytic treatment as well as these noninvasive MRI biomarkers in clinical trials in tauopathy associated neurological disorders.

## Introduction

Aging stands as the primary risk factor for the onset of Alzheimer’s disease (AD) and related dementia (1). Cellular senescence has emerged as a promising target for intervention of aging related diseases, though the underlying processes driving chronic neurodegeneration are largely unknown(2–4). Intervening in the process of biological aging and eliminating senescent cells could potentially modify the progression of these aging related disorders, offering a novel and complementary approach to existing treatment strategies(5, 6). Rodent models have played a crucial role in establishing preclinical evidence essential for translating senescence-targeting strategies to clinical applications (7). Transcriptomic comparisons between neurons with or without abnormal tau neurofibrillary tangles from postmortem human brains have unveiled a senescence signature in neurofibrillary tangles-bearing neurons(8). Thus, targeting a fundamental cellular aging process, cell senescence, presents a promising approach. Consequently, previous studies have suggested that eliminating senescent cells can enhance brain structure and function in rodent models exhibiting advanced Aβ pathology (9) and tau pathology (10, 11).

A cocktail of two US Federal Drug Administration (FDA)-approved ‘senolytic’ compounds, dasatinib and quercetin (D + Q), can selectively eliminate senescent cells from pathological tissues (12–14), and may thereby counteract age-related pathogenic processes (15, 16). Dasatinib was originally developed as an anti-cancer drug, whereas quercetin is a flavonoid found in many fruits and vegetables. Intermittent treatment with these best characterized senolytic agents, D+Q, selectively cleared senescent cells (11) which was independently reproduced in pathogenesis relevant to AD (9). Encouraged by these promising preclinical findings, the senolytic therapy strategies are progressing towards clinical testing, particularly in the context of amnestic mild cognitive impairment and early AD (NCT04063124 and NCT04685590). However, a significant challenge of clinical trials is the absence of reliable and noninvasive biomarkers to evaluate efficacy, given the inaccessibility of living human brain. Moreover, while preclinical studies have shown promising results in enhancing memory through senolytic treatment in AD models, the mechanisms on how senolytic therapy slows AD progression remain poorly understood. The knowledge gap hinders the advancement of senolytic therapies with higher efficacy and reduced side effects to clinical application.

Recent studies show that most genetic risk variants associated with late-onset AD are present in microglia, indicating that microglia may play a causal role in the disease (17–19). Chronic microglial activation can also drive pathological tau aggregation in AD (20). Microglial activation amplifies tau fibrillization in tauopathy mouse models, inflammation activates kinases that phosphorylate tau, and microglial depletion halts tau propagation(21–23). These studies point to a contributing role of microglia-driven inflammation in pathological tau and the progression of AD.

In this study, we identified noninvasive MRI measures that respond to senolytic treatment in a preclinical setting, offering potential biomarkers for future clinical trials. Our data reveals, for the first time, that water extraction with phase-contrast arterial spin tagging (WEPCAST) MRI detects impaired blood-brain barrier (BBB) integrity in a tauopathy mouse model, and chronic treatment with D+Q (weekly for 24 weeks) preserves BBB integrity in PS19 mice. Moreover, our data indicates that D+Q treatment mitigates the cerebral hypometabolism observed in PS19 mice. Collectively, these results provide compelling evidence for considering the inclusion of noninvasive MRI measures in future clinical trials, given the established MRI methods in human brains. Furthermore, this study illustrates the efficacy of D+Q senolytics in alleviating tau hyperphosphorylation, attenuating brain atrophy, and improving cognitive function in PS19 mice. Moreover, D+Q treatment shifts microglia from a disease-associated to a homeostatic phenotype, revealing a new mechanism underlying senolytic treatment. These findings underscore the importance of developing targeted therapeutic strategies to address age-related neurovascular dysfunction.

## Results

### Impaired BBB integrity and its response to D+Q treatment in 9 months old PS19 mice

BBB controls the molecular exchange between the brain parenchyma and blood, allows the selective removal of metabolic waste from the brain, plays a major role in regulating cerebral blood flow, and connects the central nervous system to the blood circulation. Recent studies have demonstrated that the compromised BBB integrity leads to the presence of pathological markers for AD (24, 25) and the extent of BBB impairment was correlated with cognitive decline (25). Therefore, the ability to investigate BBB functional integrity using non-invasive and sensitive human applicable imaging techniques has special significance in identifying biomarkers for AD. Our recently developed non-invasive MRI technique, referred to as WEPCAST, permits the assessment of BBB permeability to water molecules(26, 27). From the magnetically labeled water signal observed at the venous side, the amount of water extravasated across BBB can be measured. By comparing the WEPCAST MRI results to histological measures of tight junction proteins, we have validated WEPCAST MRI as a sensitive measure of BBB permeability (26). Using this technique, we found that patients with mild cognitive impairment manifested an increased BBB permeability to water which was correlated with the burden of amyloid and phosphorylated tau (28), suggesting a BBB breakdown occurs at the early stage of AD.

To investigate BBB integrity in tau-associated pathology, we utilized WEPCAST MRI and longitudinally assessed BBB permeability in PS19 mice at 3, 6, and 9 months (Figure 1A). Initially, there were no significant differences in BBB permeability, as determined by water extraction fraction and global cerebral blood flow, between PS19 and wild-type (WT) mice at 3 and 6 months (Figure S1). However, at 9 months old, PS19 mice displayed significantly increased water extraction fraction and BBB permeability to water compared to WT mice, while global cerebral blood flow (CBF) remained similar (Figure 1B-E). Furthermore, we conducted immunostaining of tight junction protein Claudin 5, pericyte marker CD13, vascular makers Collagen IV and CD31. We did not observe significant differences in the intensity of immunosignals of any of these proteins between wild type mice and PS19 mice at 9 months of age (Figure S2). Taken together, our results indicate that WEPCAST MRI offers a more sensitive measure of early BBB permeability disruption compared to traditional immunostaining of BBB structural protein changes.

**Figure 1.**
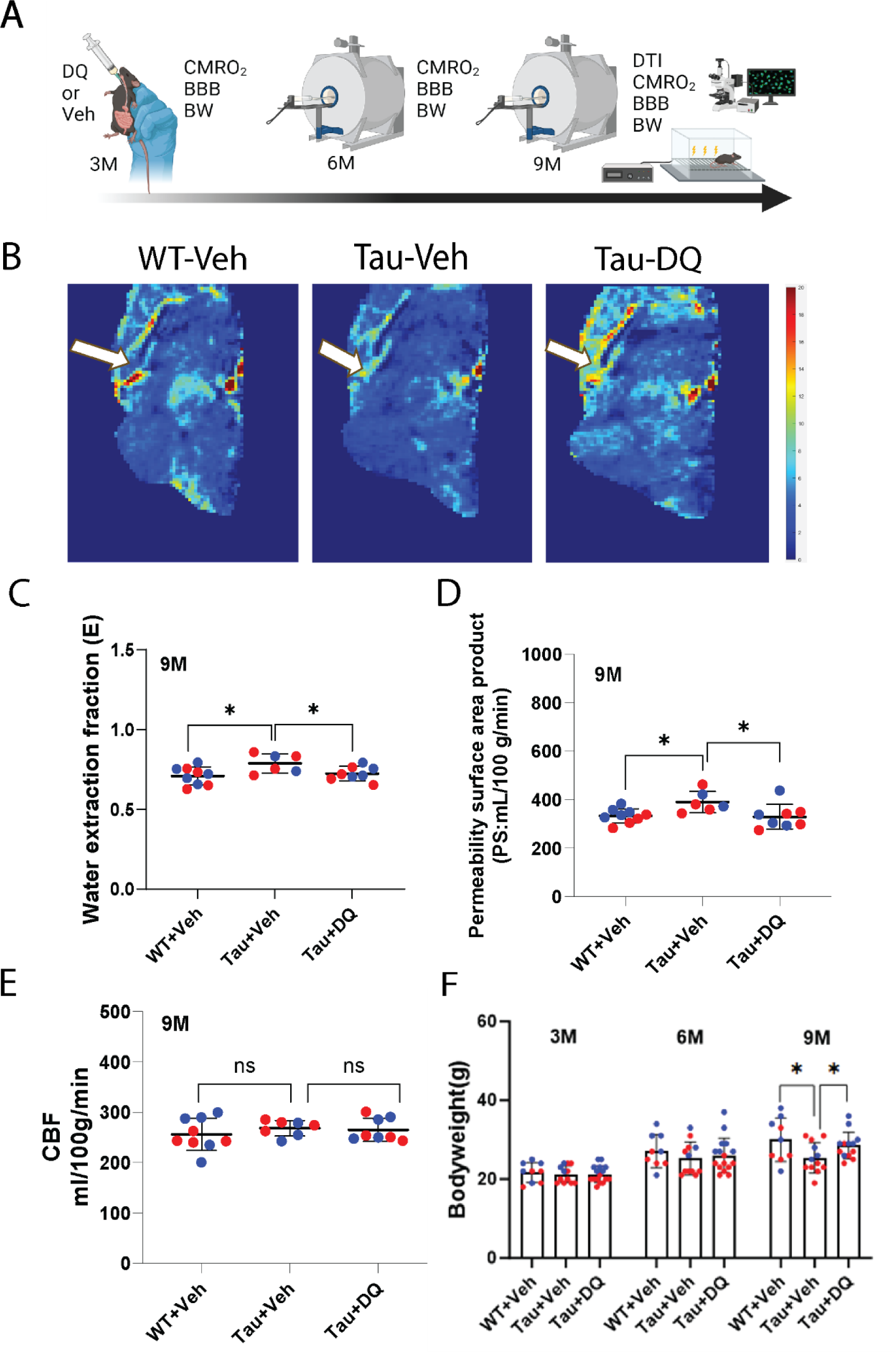
DQ treatment maintains blood-brain barrier (BBB) integrity in 9 months old PS19 mice. (**A**) Timeline of the experiment design and outcome measurements. (**B**) Representative images of WEPCAST MRI in indicated groups. Arrows indicate the Great vein of Galen (GVG) where the WEPCAST MRI detects the amount of magnetically labelled water. Warm color indicates higher amounts of magnetically labelled water. Note that PS19 brain has reduced magnetically labelled water in the GVG. (**C** and **D**) Water extraction fraction and permeability surface area product (PS) were calculated according to the formulas described in the method section. Higher water extraction fraction and PS values indicate increased BBB permeability to water. n=6-9 mice/group *p<0.05 by one-way ANOVA with Bonferroni correction for multiple comparisons. (**E**) Cerebral blood flow (CBF) evaluated by MRI in the same mice was used for BBB measurement. n = 7-9 mice/group; ns, no statistical significance by one-way *ANOVA* with Bonferroni correction for multiple comparisons. (**F**) Body weights of the mice underwent BBB and CBF measures at indicated ages. n = 9-15 mice/group; *p<0.05 by one-way ANOVA with Bonferroni correction for multiple comparisons. In all graphs, each individual data points are indicated; Red dots represent female mice and blue dots represent male mice. All data are shown as individual data points, Mean and SD. Wild type treated with vehicle (WT+Veh), PS19 mice treated with vehicle (Tau +Veh) or DQ (Tau+DQ).

To further investigate whether WEPCAST BBB permeability measures respond to treatment, we evaluated D+Q senolytic therapy, known for its protective effects against AD in tauopathy models. We implemented a chronic treatment schedule starting at 3 months, where D (5mg/kg) + Q (50 mg/kg) or vehicle (20% PEG) were administered orally at a weekly basis until the end of the study. Notably, D+Q treatment preserved BBB integrity in 9-month-old PS19 mice, restoring it to the normal range observed in wild-type littermate controls (Figure 1B-D), without impacting CBF (Figure 1E). Remarkably, PS19 mice in the vehicle-treated group exhibited significant weight loss at 9 months old, while D+Q treatment rescued their body weight (Figure 1F).

### D+Q treatment mitigates hypometabolism and attenuates brain atrophy in 9 months old PS19 mice

Cerebral hypometabolism, a characteristic feature observed earlier in human Alzheimer’s disease (AD) before neuropathology and symptoms manifest, continues to worsen as symptoms progress, and is more pronounced than that of normal aging (29). To assess whether cerebral metabolism is altered in PS19 mice, we utilized T2 relaxation under spin tagging (TRUST) MRI to longitudinally evaluate cerebral metabolism(30), as indicated by oxygen extraction fraction (OEF) and cerebral metabolic rate of oxygen (CMRO2), in mice aged 3 (Figure SA-B), 6, and 9 months old. Our findings indicate that PS19 mice begin to exhibit hypometabolism at 6 months of age, manifested by reduced OEF (Figure S2C) and CMRO2 (Figure S2D). Starting at 3 months old, PS19 mice were randomly assigned to receive either vehicle (20% PEG) or D+Q treatment weekly for 6 months. Interestingly, D+Q treatment shows a trend towards improving the hypometabolic status of PS19 mice at 6 months of age, following 3 months of D+Q treatment (Figure S2 C-D). At 9 months old, PS19 mice continued to display reduced OEF (Figure 2A) and a declining trend in CMRO2 (Figure 2B). In contrast, D+Q pretreatment prevented the cerebral metabolism deficits in PS19 mice, as indicated by normalized OEF and CMRO2 (Figure 2A-B).

**Figure 2.**
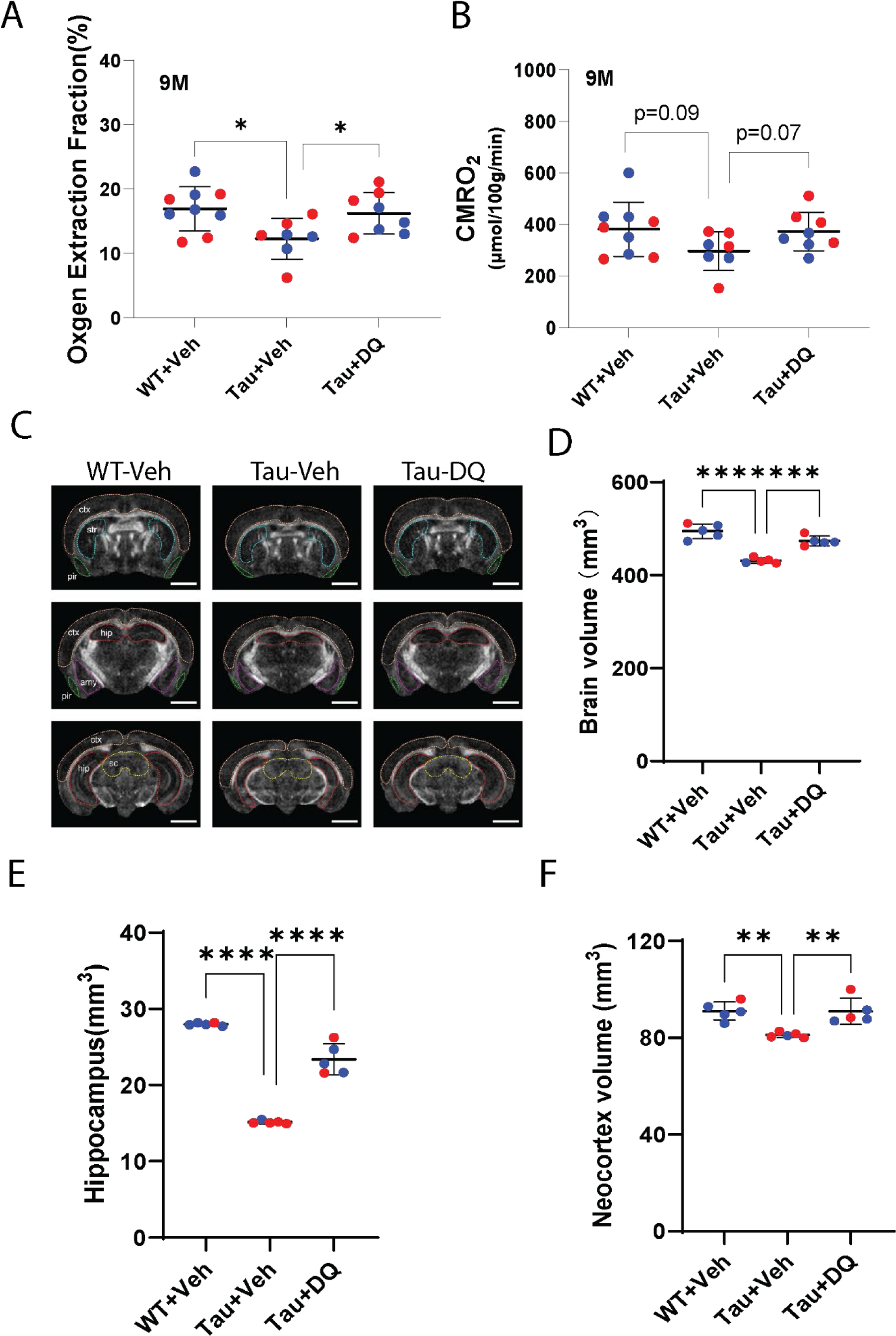
DQ treatment mitigates hypometabolism and attenuates brain atrophy in 9 months old PS19 mice. (**A** and **B**) Oxygen extraction fraction (OEF) and global cerebral metabolic rate of oxygen (CMRO2) were detected by noninvasive T2 relaxation under spin tagging (TRUST) and phase contrast MRI in vivo. n = 7-9 mice/group; *p<0.05 by one-way ANOVA with Tukey post hoc analysis. (**C**) Representative images of brain volumetric MRI, ctx, Cortex; str, Striatum; hip, hippocampus; pir, piriform cortex. Scale bar = 2mm. (**D-F**) Brain volumes were measured by high resolution diffusion tensor MRI at the indicated brain regions. n=5 mice/group; **p<0.01; ****p<0.0001 by *ANOVA* with Bonferroni correction for multiple comparisons. In all graphs, each individual data points are indicated; Red dots represent female mice and blue dots represent male mice. All data are shown as individual data points, Mean and SD. Wild type treated with vehicle (WT+Veh), PS19 mice treated with vehicle (Tau +Veh) or DQ (Tau+DQ).

Brain atrophy, a pathological hallmark of AD, akin to human patients, was evaluated using high-resolution Diffusion tensor imaging (DTI) MRI to measure subregional brain volumes. As shown in prior work (22), PS19 mice exhibited significant brain atrophy in multiple brain regions at 9 months of age (Figure 2C-D), particularly in the hippocampus (Figure 2E) and neocortex (Figure 2F). Remarkably, treatment with D+Q (weekly for 6 months) significantly preserved brain volumes towards the range observed in wild-type mice (Figure 2C-F), indicating that chronic treatment with D+Q attenuates brain atrophy in PS19 mice.

### D+Q treatment reduces tau hyperphosphorylation and improves cognitive function in 9 months old PS19 mice

A distinguishing characteristic of PS19 mice is the development of protein aggregates consisting of hyperphosphorylated tau protein, identifiable by phospho-tau (Ser 202, Thr 205) monoclonal antibody (AT8). To assess the impact of D+Q senolytic treatment on tau hyperphosphorylation, we evaluated AT8 positive signals, indicative of phosphorylated Tau levels. As shown in prior work (22), vehicle-treated PS19 mice exhibited abundant AT8 signals in neurons (Figure 3A). Conversely, D+Q-treated PS19 mice showed markedly fewer AT8-positive neurons, suggesting reduced phosphorylated tau levels (Figure 3A,B). These findings suggest that the elimination of senescent cells ameliorates tau hyperphosphorylation and prevents tau aggregations.

**Figure 3.**
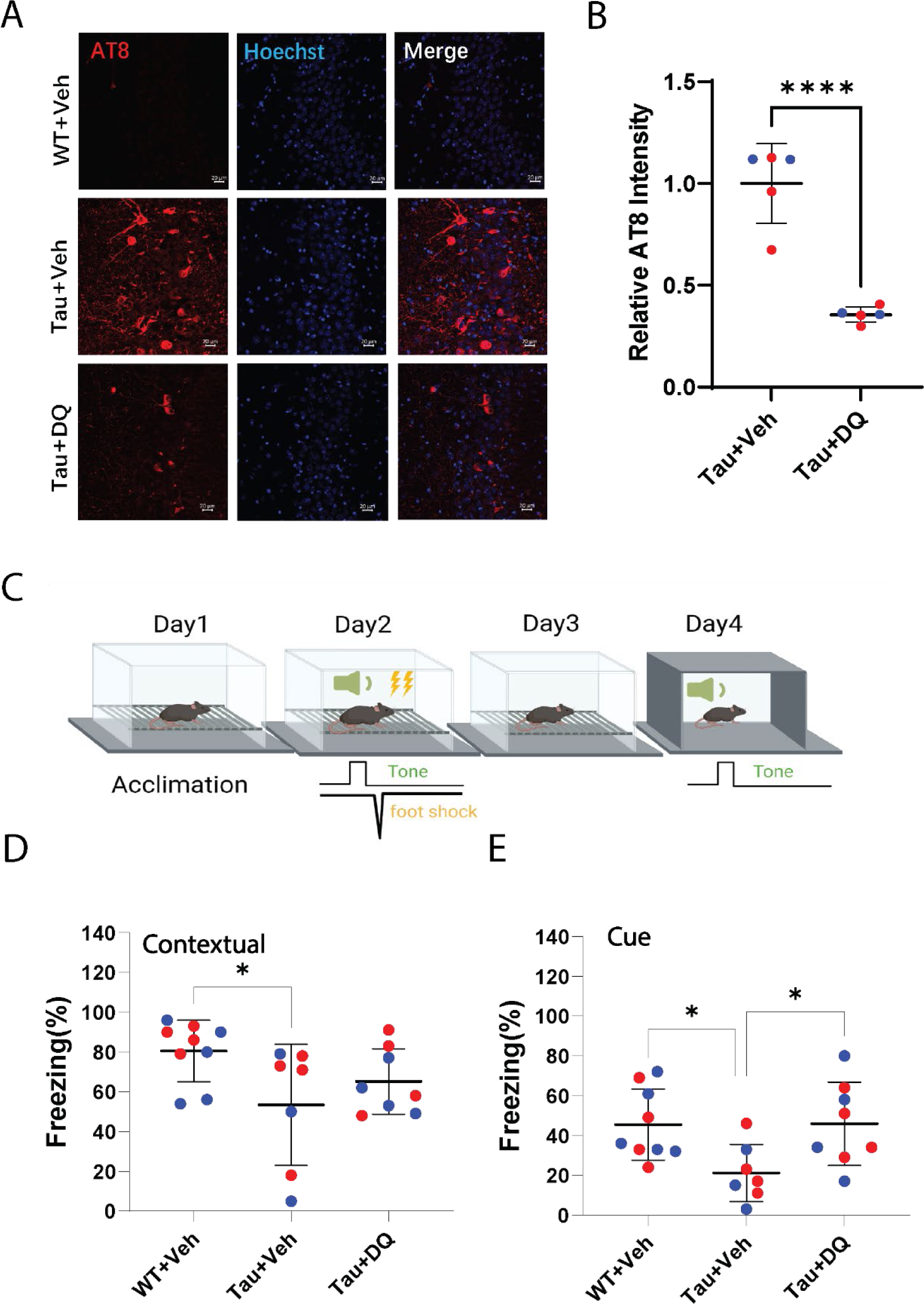
DQ treatment reduces tauopathy and improves learning deficits in the trace fear conditioning test of 9 months old PS19 mice. (**A**) Representative images of phosphorylated Tau (AT8 staining) in the CA3 of hippocampus in the indicated groups. Scale bar = 20 µm. (**B**) Quantification of phosphorylated Tau levels in PS19 mice treated with vehicle (Tau +Veh) or DQ (Tau+DQ). N=5 mice/group; ****p<0.0001 by unpaired Student’s t-test. (**C**) Schematic illustration of the trace fear conditioning (TFC) test. (**D** and **E**) Comparison of learning capacity by freezing response to contextual (D) or cued (E) stimulus. n=7-9 mice/group; *p<0.05 by *ANOVA* with Bonferroni correction for multiple comparisons. In all graphs, each individual data points are indicated; Red dots represent female mice and blue dots represent male mice. All data are shown as individual data points, Mean and SD. Wild type treated with vehicle (WT+Veh), PS19 mice treated with vehicle (Tau +Veh) or DQ (Tau+DQ).

To further investigate whether the observed beneficial effects of D+Q administration improved cognitive function, we conducted the trace fear conditioning test. This test involves a neutral conditioned stimulus (tone) and an aversive unconditioned stimulus (foot shock), separated by a trace interval (Figure 3C), which engages the hippocampus. Mice were exposed to five pairings of tone and foot shock with a 20-second trace interval. Contextual memory was assessed 24 hours after conditioning by measuring freezing behavior when placed in the conditioned chamber. Additionally, 48 hours after conditioning, freezing behavior was measured in a dark chamber with only the tone stimulus, serving as a cue.

We found that vehicle-treated PS19 mice exhibited deficits in both contextual and cue-dependent memory, indicative of hippocampal-dependent memory impairment (Figure 3D,E). In contrast, D+Q-treated PS19 mice displayed increased freezing events in the conditional chamber compared to vehicle-treated PS19 mice (Figure 3D), suggesting better-preserved contextual memory. Furthermore, D+Q-treated PS19 mice behaved similarly to littermate control mice in the cue-dependent memory test (Figure 3E). These results collectively demonstrate that senescent cells may contribute to the loss of hippocampus-dependent learning and memory in PS19 mice, and senolytic therapy reduces tau hyperphosphorylation while improving cognitive function.

### D+Q treatment shifts microglia from a disease-associated to a homeostatic phenotype

To comprehend the mechanistic contribution of senescence to tau-mediated pathology, we aimed to identify the specific cell types undergoing senescence. The protein p16/INK4a serves as a cell cycle inhibitor and is associated with cell senescence, commonly used as a marker for labeling senescent cells. Brain sections were co-stained with p16/INK4a and microglia markers IBA1. Notably, p16/INK4a immunoreactive cells were not observed in the hippocampus of wild-type mice, serving as a control (Figure 4A,). However, a substantial increase in p16/INK4a immunoreactivity was observed in IBA1-positive microglia in vehicle-treated PS19 mice compared to wild-type mice (Figure 4A-B). Conversely, D+Q-treated PS19 mice exhibited a marked reduction in p16/INK4a immunoreactivity in IBA1-positive microglia (Figure 4A-B).

**Figure 4.**
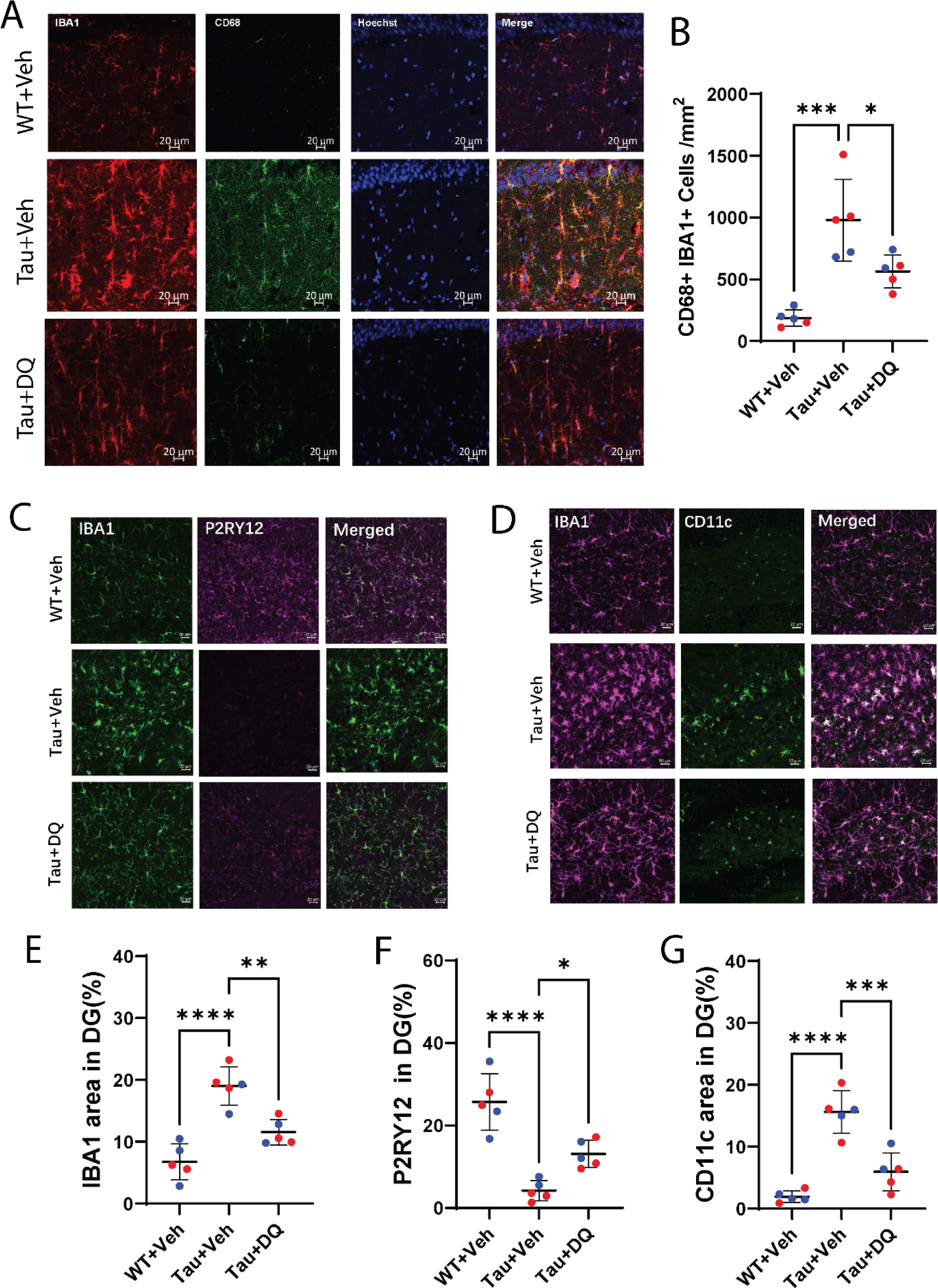
DQ treatment shifts microglia from a disease-associated subtype to a homeostatic subtype. (**A)** Representative images of activated microglia (CD68 positive microglia) in the hippocampus. Scale bar =20 µm. (**B**) Quantification of activated microglia in the hippocampus at indicated groups. n= 5; \**p*<0.05, \*\*\**p*<0.001 by ANOVA with Bonferroni correction for multiple comparisons. (**C**) Representative images of microglia marker IBA1 and homeostatic microglia marker P2RY12 staining in the hippocampus (dentate gyrus). Scale bar = 20 µm. (**D**) Representative images of microglia marker IBA1 and disease-associated microglia marker CD11c staining in the hippocampus (dentate gyrus). Scale bar = 20 µm. (**E**-**G**) Quantification of IBA1, P2RY12, and CD11c staining in the hippocampus of mice in the indicated groups. n= 5 mice/group; *p<0.05, **p<0.01, ***p<0.001, ****p<0.0001 by *ANOVA* with Bonferroni correction for multiple comparisons. In all graphs, each individual data points are indicated; Red dots represent female mice and blue dots represent male mice. All data are shown as individual data points, Mean and SD. Wild type treated with vehicle (WT+Veh), PS19 mice treated with vehicle (Tau +Veh) or DQ (Tau+DQ).

Microglia expresses a range of surface receptors enabling them to detect environmental changes. IBA1 serves as a pan-microglia marker. Vehicle-treated PS19 mice displayed significantly increased IBA1-positive cells, indicative of microgliosis in these mouse brains (Figure 4A-C). Conversely, D+Q treatment significantly reduced the numbers of IBA1-positive microglia in PS19 mice (Figure 4A-C). Across species and models, diverse and context-dependent microglial states have been observed. Notably, disease-associated microglia (DAMs) have been associated with AD pathology models. An upregulation of CD11c and downregulation of P2RY12 have been indicative of a microglial state associated with aging or neurodegenerative disease models. P2RY12 serves as a homeostatic marker for microglia.

Given the senescent phenotype exhibited by microglia in the tau pathology environment, we investigated whether microglial states were associated with senescence markers in the brains of PS19 mice. In the vehicle-treated wild-type control group, the majority of IBA1 immunoreactive microglia expressed high levels of P2RY12 immunoreactivity but low CD11c immunoreactivity. To elucidate the influence of D+Q treatment on microglial states, we conducted double immunostaining with IBA1 antibody combined with either the well-characterized DAM marker CD11c or the homeostatic marker P2RY12.

Although overall microglial numbers were increased in PS19 mice treated with vehicle compared to wild-type mice, P2RY12-positive microglia were significantly fewer in vehicle-treated PS19 mice, while more CD11c-expressing microglia were observed. However, PS19 mice treated with D+Q (weekly for 24 weeks) exhibited significantly more P2RY12 staining microglia (Figure 4A,D) and fewer CD11c-positive microglia (Figure 4B,E). These findings collectively suggest that senolytic therapy with D+Q not only alleviates microgliosis but also promotes microglial homeostasis by shifting microglia from a DAM to a homeostatic state.

## Discussion

Emerging evidence suggests that tau protein accumulation drives cellular senescence in the brain. Pharmacologically clearing senescent cells in mouse models of tauopathy reduced brain pathogenesis (10, 11). Our findings herein provide validating support for prior work demonstrating benefits of D+Q treatment using MRI as outcome measures and indicate that compared to vehicle-treated tauopathy mice (11, 31), intermittent senolytic therapy with D+Q reduced tau accumulation, preserved BBB integrity and cerebral metabolism, attenuated brain atrophy, sustained microglia homeostasis by reducing disease associate microglia subtype and promoting its homeostatic phenotype, and improved cognitive performance in a tauopathy mouse model. Similar to previous structural MRI studies, our study also demonstrated that D+Q therapy preserves brain tissue of AD mice. Furthermore, our noninvasive functional MRI measures of BBB permeability and CMRO2 have expanded our understanding of the metabolic outcomes associated with D+Q treatment. Noninvasive imaging outcomes are crucial parameters in clinical studies, as demonstrated in the Phase 2 D+Q trial in AD (SToMP-AD, NCT04685590). Our data underscores the utility of noninvasive imaging methods for evaluation, highlighting their power and reliability in anti-senolytic studies (31).

Intermittent dosing of the D+Q has shown an acceptable safety profile in clinical studies for other senescence-associated conditions(32). This study further provides strong proof-of-concept evidence that tau-induced cellular senescence is a novel therapeutic target for neurodegenerative diseases characterized by abnormal tau deposition in the brain (10, 11). By incorporating noninvasive MRI measures in this preclinical study, we identified that MRI measures of the BBB permeability, cerebral metabolism, and brain volumetric changes could be considered in the future clinical trials of evaluating treatment efficacy.

Our data indicates a significant decrease in the cerebral metabolic rate of oxygen in the PS19 tauopathy model, while the global cerebral blood flow shows no significant difference. Previous studies suggest that tau deposition leads to senescence-associated transcriptomic changes (10, 11, 33), followed by vascular alterations in the mouse brain, such as transient capillary occlusion. (33). These vascular changes can disrupt capillary transit and impair neuronal oxygenation without gross reductions in cerebral blood flow. Our data further support this notion in the PS19 tauopathy model. Additionally, we observed decreased cerebral metabolic rate of oxygen in 6-month-old PS19 mice when blood-brain barrier permeability is still intact, suggesting that these brain metabolic changes are early pathological events.

WEPCAST MRI quantifies the relative fraction of magnetically labeled spins that penetrate brain tissue versus those remaining in veins. Notably, water spins from blood enter brain tissue through the BBB. Increased water spins in arterial blood indicate “leakage” into brain tissue upon BBB disruption. Quantitatively, the fraction of water spins entering brain tissue is termed the water extraction fraction (E). Using the Renkin-Crone equation, the permeability-surface-area product (PS) can be derived as an index of BBB integrity.

The WEPCAST technique benefits from high sensitivity by employing small water molecules (18 Da) as tracers, unlike fluorescent tracers used for parenchymal leakage detection. This method has been validated in a mouse model of Huntington’s disease (26) and in human studies with Alzheimer’s disease patients (28), demonstrating its efficacy in detecting BBB disruption.

In WEPCAST MRI images, increased arterial blood spins entering brain tissue due to BBB leakage correspond to decreased magnetically labeled spins detected from the venous side, resulting in a less bright signal at the Great Vein of Galen region. A higher quantified PS value indicates greater BBB permeability, indicative of increased leakage.

Isoflurane is recognized as a modulator of the blood-brain barrier (BBB) in a dose-dependent manner (34). Literature has compared BBB function under different isoflurane concentrations (1.0% vs. 3.0%), indicating that significant BBB opening primarily occurs at higher doses. In our experiments, we consistently employed low doses of isoflurane for both induction and maintenance. Therefore, any observed changes in permeability were minimally influenced by the anesthetic agent and predominantly reflected pathological and treatment effects.

The blood-brain barrier, a crucial component of the neurovascular unit (35), represents a highly dynamic system where the proper functioning of the brain relies on the functional interactions of endothelial cells, neurons, pericytes, and glial cells. The BBB strictly regulates the exchange of cells and molecules between blood and brain parenchyma. It has been reported that disruption of the BBB found in tauopathies is driven by chronic neuroinflammation (36). The neuropathological hallmarks such as intra and extracellular protein aggregates are associated with chronic neuroinflammation has long been known. Chronic neuroinflammation affects the BBB integrity by increasing vascular permeability, promoting structural changes in brain capillaries such as fragmentation (37, 38). Inflammatory mediators play a pivotal role in regulating blood-to-brain cell transmigration, perpetuating inflammation, and exacerbating disease pathology. Structural and functional changes in the BBB can result in progressive synaptic and neuronal dysfunction (35, 39). Our data indicate that BBB permeability is impaired when microgliosis is evident in the brains of PS19 mice. More importantly, chronic treatment with D+Q preserved BBB integrity and attenuated microgliosis. This is the first demonstration that a noninvasive MRI measure of BBB permeability responds to therapeutic agents in a tauopathy mouse model. These findings lay the groundwork for considering the future application of MRI measures of BBB permeability in clinical trials involving senolytic therapy and other potential disease-modifying treatments for tauopathy-related neurological disorders.

Regarding body weight, there was no significant difference between 3-month-old and 6-month-old wild-type and PS19 mice. However, PS19 mice had lower body weight at 9 months, which may be related to pathological progress. D+Q therapy partially reversed this weight loss, suggesting that senolytic treatment not only impacts tau deposition and disease-related neurological disorders, but also improves general health including food intake and nutrition absorption.

It has been shown that tau tangle formation correlated with neuroinflammation in AD (40). Moreover, neuroinflammation linked to tau deposition has been well documented in mice transgenic for human mutant tau protein (22, 41). Microglia are the major cell type contributing to neuroinflammation. It is now clear that microglia exist in diverse, dynamic, and multidimensional stages depending on the context, including local environment. Microglia have a complex ‘‘sensome’’(42) and a series of surface receptors that allow them to detect changes in their environment. One subtype of microglia, disease-associated microglia (DAMs), were initially identified in a mouse model of AD that expresses five human familiar AD mutations (5XFAD) and further validated in other amyloid-β AD mouse models (43), and then evident across several disease models (44). There are specific marker proteins to distinguish the subtype of microglia. CD11c is mainly restricted to the plasma membrane and highly expressed in DAMs and often is used to identify DAMs. While P2RY12 localizes to the plasma membrane or diffuses throughout the cytoplasm, it is preferentially expressed in homeostatic microglia.

Our data indicates that increased overall microglia populations in the PS19 mouse brain, particularly in the hippocampus. Similarly, a previous study by Brelstaff et al. showed that microglia undergo senescence following the phagocytosis of neurons containing pathogenic tau. Their findings provide additional validation that microglia enter a senescent state in PS19 mice, supporting our study (45). Among the overall increased microglia in the PS 19 mouse hippocampus, there is higher ratio of DAMs subtype than that of homeostatic microglia compared to control mice. Interestingly, senolytic treatment with D+Q shifted the microglial phenotype from DAMs to a homeostatic subtype, while also reducing overall microgliosis. These observations are particularly significant, as microglia are highly responsive to neurodegenerative conditions, undergoing changes in their molecular profile, motility, and function.

The expression of core microglial markers is also altered over the course of disease. Downregulation of the homeostatic microglial signature, P2RY12, is one of the salient features of the microglial response to AD pathology, as shown in several mouse models of disease (46). Chronic D+Q treatment can preserve microglia in its homeostatic feature.

The mammalian brain is among the most metabolically active organs in the body and requires large amounts of energy for proper function. The high energy demands require efficient oxidative metabolism support. Besides the classical pathology of AD, which is characterized by cerebral atrophy, with senile plaques, dystrophic neurites, and neurofibrillary tangles within defined areas of the brain. Another characteristic of AD is regional hypometabolism in the brain. This decline in cerebral glucose metabolism occurs before pathology and symptoms manifest, continues as symptoms progress, and is more severe than that of normal aging. Glucose hypometabolism appears to manifest in at-risk individuals decades before clinical symptoms of dementia emerge, and it is unlikely to be solely attributed to cell loss. Considering that hypometabolism is an early and progressive event in AD and may trigger downstream pathological events, targeting this process as a treatment for AD is reasonable. Therefore, a therapeutic goal in AD and other cognitive disorders is to enhance the neuronal energy state. Our results suggest that senolytic treatment with D+Q showed a trend towards improved oxygen extraction fraction and CMRO2.

In conclusion, our findings suggest that targeting cellular senescence effectively mitigates a chronic neurodegenerative cascade, reducing tau hyperphosphorylation-associated neuroinflammation, preserving BBB integrity, and ameliorating brain atrophy and hypermetabolism, ultimately leading to improved cognitive function. Overall, our data provide evidence that cellular senescence is a fundamental pathogenic process common among AD pathologies, reaffirming that cell senescence is a druggable target for treating various tau-associated neurodegenerative diseases. Furthermore, our results pave the way for future application of MRI measures of BBB permeability and cerebral metabolism in clinical trials, as the MRI methods employed in this study are ready and implementable for human studies at clinical scanners (27, 47).

## Materials and Methods

### Animals and brain samples

Transgenic mice. Male and female PS19 transgenic mice (B6;C3-Tg(Prnp-MAPT*P301S)PS19Vle/J stock no. 008169), and age-matched wild-type controls, were obtained from the Jackson Laboratories. Mice were housed in sex-matched groups of five in standard mouse cages on a 12-h light/dark cycle at a room temperature (RT) of 23◦C with free access to food and water. Mice were eventually anesthetized with isoflurane and sequentially perfused transcardially with phosphate-buffered saline (PBS) and 4% paraformaldehyde in PBS. Fixed and sucrose-cryoprotected brain tissue was cut horizontally with a cryostat (Leica, CM1950) into 40-μm sections. Fresh frozen brains were also carefully collected after anesthesia with isoflurane. All procedures were approved by the Animal Care and Use Committee of the Johns Hopkins University, School of Medicine.

### DQ treatment regimen

3-month-old PS19 mice were randomly assigned to treatments by oral gavage with dasatinib (Selleckchem) and quercetin (Selleckchem) or vehicle once weekly for 24 weeks. Dasatinib and quercetin were first dissolved in pure polyethylene glycol (PEG, Sigma-Aldrich) by sonication for 5min, at a final concentration of 12mg/kg of dasatinib and 50mg kg–1 of quercetin in 20% PEG with 0.9% saline for treatment. The mice in vehicle group were treated with 20% PEG in 0.9% saline. Body weights of mice in each group were recorded every week before dose calculating and gavage. MRI scans were conducted 3 days after the last DQ administration, and behavioral tests were conducted 5 days after the last DQ administration.

### Trace Fear Conditioning (TFC) test

TFC was conducted at 9 months old. Briefly, TFC consisted of a habituation day, and three consecutive training and testing days. On the habituation day, mice were exposed to the shock box (Coulbourn, Holliston, MA, USA) for 10 min. On the training day, mice were placed in the shock box and given a 2-min habituation, after which a 20-second white noise tone (80 db, 2000 Hz) was delivered. Twenty seconds following the termination of the tone, a scrambled 2-s 0.5 mA shock was delivered. The tone-shock pairing was repeated three additional times. On the test days, mice were placed in the shock box for 3 min to measure freezing in response to the context. Mice were then placed in a separate context and freezing in response to the 20-second white noise. Freezing behavior was automatically scored using Cleversys Freezescan (Cleversys Inc., Reston, VA, USA).

### Immunohistochemistry and quantification

Staining was performed on free-floating mouse brain sections. The sections were rinsed three times for 10min in PBS and then were incubated with blocking buffer (5% goat or donkey serum and 0.2% Triton X-100 in PBS) for 1h at room temperature. Then the sections were incubated with primary antibodies in blocking buffer overnight at 4 °C: AT8 (Thermo Scientific, MN1020, Mouse,1:100), IBA1(Wako, 019-19741, Rabbit, 1:1000, or Abcam, ab283346 Rat 1:100), CD68 (Biorad, MCA1957, Rat, 1:100), CD11C (Cell Signaling Technology, 975855, Rabbit, 1:100), P2RY12(Biolegend, 848002, Rat 1:100), p16/INK4a (Abcam, ab 54210, mouse, 1:50). After thorough washes in PBS, sections were incubated with 1:1000 dilution of Alexa 488-, Alexa 555-, and Alexa 647-conjugated secondary antibodies (Thermo Scientific, Goat anti-rabbit 488: **A11008**, Goat anti-mouse 488: **A11001,** Goat anti-rabbit 555: **A32732**, Goat anti-mouse 555: **A21422**, Goat anti-rat 647: **A21247**) appropriate for the species of the primary antibodies at room temperature for 1 h, followed by Hoechst counterstains. The sections were examined and images acquired using Zeiss LSM700 laser-scanning confocal microscopes. For quantitation of Tauopathy and fluorescence intensities of IBA1, CD11c and P2RY12 in hippocampus, images of CA3 and dentate gyrus areas were acquired using a Zeiss LSM700 microscope with a ×20 objective. Analyses were performed using Fiji ImageJ software. For quantification, five consecutive sections per mouse were used, with five mice in each group. From each section, four images were selected for quantification. Mean intensity values were calculated from these images to obtain final results for statistical analysis.

### Functional MRI

Functional MRI were conducted at an 11.7T Bruker Biospec system (Bruker, Ettlingen, Germany) with a horizontal bore equipped with an actively shielded pulse field gradient (maximum intensity of 0.74 T/m). Images were acquired using a 72-mm quadrature volume resonator as a transmitter, and a four-element (2×2) phased-array coil as a receiver. The homogeneity of B0 field over the mouse brain was optimized with a global shimming (up to 2nd order) based on a subject-specific pre-acquired field map.

Functional MR imaging was conducted under low-dose isoflurane anesthesia carried by medical air (21% O2, 78% N2). Anesthesia induction was at 1.5% concentration for 15 minutes, followed by maintenance at 1.0%. The respiration rate of each mouse was continuously monitored during experiments to ensure survival and maintain consistent respiratory rates across all experimental mice. If a mouse exhibited a breathing rate exceeding 150 breaths per minute, the maintenance isoflurane dose was slightly increased to 1.2%. This anesthesia protocol has been previously utilized and documented (48).

Additionally, each mouse was immobilized using a bite bar and ear pins, then placed on a water-heated animal bed with temperature control.

*Oxygen extraction fraction (OEF)* is defined as the arteriovenous difference in blood oxygenation and was measured by the T2-relaxation-under-spin-tagging (TRUST) MRI technique. TRUST scan was implemented following the reported MRI protocol(30). Key parameters were: TR/TE = 3500/6.5 ms, FOV = 16×16 mm^2^, matrix size = 128×128, slice thickness = 0.5 mm, inversion-slab thickness = 2.5 mm, post-labeling delay = 1000 ms, eTE = 0.25, 20, 40 ms, and scan duration = 5.6 min with two repetitions.

Cerebral blood flow (CBF) was evaluated with phase contrast (PC) MRI focusing on the three major feeding arteries of brain (left/right internal carotid arteries and basilar artery). The previously reported protocol was utilized(49), and the key parameters were: TR/TE = 15/3.2 ms, FOV = 15×15 mm^2^, matrix size = 300×300, slice thickness = 0.5 mm, receiver bandwidth = 100 kHz, flip angle = 25°, and scan duration = 0.4 min per artery. Brain volume was estimated from a T_2_-weighted fast-spin-echo MRI protocol (TR/TE = 4000/26 ms, FOV = 15×15 mm^2^, matrix size = 128×128, slice thickness = 0.5 mm, 35 axial slices, and scan duration = 1.1 min)(30).

*Blood-brain barrier* function was assessed with WEPCAST MRI(25). Key parameters were: TR/TE = 3000/11.8 ms, labeling duration = 1200 ms, FOV = 15×15 mm^2^, matrix size = 96×96, slice thickness = 1 mm, labeling-pulse width = 0.4 ms, inter-labeling-pulse delay = 0.8 ms, flip angle of labeling pulse = 40°, and scan duration = 4.0 min with two-segment spin-echo echo-planar-imaging acquisition.

### Structural MRI and DTI

Ex vivo structural MRI was performed on a vertical-bore 11.7 T MRI scanner (Bruker Biospin) equipped with an actively-shielded Micro2.5 gradient system (maximum intensity of 1.5 T/m). The mouse skulls were placed in 20-mm-diameter tubes that were filled with Fomblin® (Solvay Inc., Princeton, NJ, USA) for susceptibility matching. A 20-mm-diameter volume coil was used as the radiofrequency transmitter and receptor. Data were acquired using a 3D diffusion-weighted gradient-and-spin-echo sequence with bipolar diffusion gradients and twin navigator echoes (50). The imaging parameters were as follows: turbo factor = 4, echo-planar-imaging factor = 3, TR/TE = 700/28 ms, FOV = 12 x 9.2 x 17.4 mm, matrix size = 120 x 92 x 174, spatial resolution = 100 µm isotropic, 2 signal averages, and receiver bandwidth = 90 kHz. Images along 30 uniform diffusion-encoding directions and 3 non-diffusion-weighted images were acquired, with a total scan time of 17 h.

Images were reconstructed from the raw *k*-space data using in-house written code in IDL (ITT Visual Information Solutions, Boulder, CO, USA). Diffusion tensors were fit to the log-signal using Mrtrix3 (51) and maps of fractional anisotropy (FA) were calculated from the tensor eigenvalues. The brain volumes were extracted using seeded region-growing segmentation of the averaged diffusion-weighted images in ROIEditor (www.mristudio.org). For volumetric analysis, the 3D brain images were deformably registered to a mouse brain DTI atlas (52). Regions of interest from the DTI atlas were warped to the native image space of each brain for atlas-based segmentation of gray matter structures including the hippocampus and neocortex.

### Statistical Analysis

Data are expressed as the individual value unless otherwise noted. Statistical analysis was performed with SPSS using two-tailed Student’s t-test, one-way ANOVA, two-way ANOVA, with Bonferroni post-hoc tests. The p-values less than 0.05 were considered statistically significant. N is reported in the figure legends.

## Data and materials availability

All data associated with this study are present in the paper or the Supplementary Materials. Additional requests for raw and analyzed data or materials should be made to and will be promptly reviewed to determine whether the application is subject to any intellectual property or confidentiality requirements. Any data and materials that can be shared will be released after the execution of a material transfer agreement.

## Acknowledgements

The study is supported by NIH R01 NS124084, NIH R01 NS127344 (to W.D); NIH R21 NS119960, NIH R01 AG081932 to (Z.W), and R01AG071515 (to H.L.).

## Author contributions

Conceptualization W.D., M.Y.; Methodology W.D., M.Y., Z.W. Experimental Work M.Y., Z.W., J.S.N., A.K., Y.O., R.L.,T.C.; Writing-original Draft. W.D., M.Y., Z.L.; Writing -Review & Editing, W.D., Z.W., S.S., H.L., M.A., Funding Acquisition, W.D., Z.W., H.L.; Supervision, W.D..

## References

1. Garcia MJ, Leadley R, Ross J, Bozeat S, Redhead G, Hansson O, et al. Prognostic and Predictive Factors in Early Alzheimer’s Disease: A Systematic Review. J Alzheimers Dis Rep. 2024;8(1):203–40.

2. de Luzy IR, Lee MK, Mobley WC, and Studer L. Lessons from inducible pluripotent stem cell models on neuronal senescence in aging and neurodegeneration. Nat Aging. 2024.

3. Richardson M, and Richardson DR. Pharmacological Targeting of Senescence with Senolytics as a New Therapeutic Strategy for Neurodegeneration. Mol Pharmacol. 2024;105(2):64–74.

4. Melo Dos Santos LS, Trombetta-Lima M, Eggen B, and Demaria M. Cellular senescence in brain aging and neurodegeneration. Ageing Res Rev. 2024;93:102141.

5. Chou SM, Yen YH, Yuan F, Zhang SC, and Chong CM. Neuronal Senescence in the Aged Brain. Aging Dis. 2023;14(5):1618–32.

6. Ma Y, and Farny NG. Connecting the dots: Neuronal senescence, stress granules, and neurodegeneration. Gene. 2023;871:147437.

7. Tchkonia T, and Kirkland JL. Aging, Cell Senescence, and Chronic Disease: Emerging Therapeutic Strategies. JAMA. 2018;320(13):1319–20.

8. Dehkordi SK, Walker J, Sah E, Bennett E, Atrian F, Frost B, et al. Profiling senescent cells in human brains reveals neurons with CDKN2D/p19 and tau neuropathology. Nat Aging. 2021;1(12):1107–16.

9. Zhang P, Kishimoto Y, Grammatikakis I, Gottimukkala K, Cutler RG, Zhang S, et al. Senolytic therapy alleviates Abeta-associated oligodendrocyte progenitor cell senescence and cognitive deficits in an Alzheimer’s disease model. Nat Neurosci. 2019;22(5):719–28.

10. Bussian TJ, Aziz A, Meyer CF, Swenson BL, van Deursen JM, and Baker DJ. Clearance of senescent glial cells prevents tau-dependent pathology and cognitive decline. Nature. 2018;562(7728):578–82.

11. Musi N, Valentine JM, Sickora KR, Baeuerle E, Thompson CS, Shen Q, et al. Tau protein aggregation is associated with cellular senescence in the brain. Aging Cell. 2018;17(6):e12840.

12. Xu M, Pirtskhalava T, Farr JN, Weigand BM, Palmer AK, Weivoda MM, et al. Senolytics improve physical function and increase lifespan in old age. Nat Med. 2018;24(8):1246–56.

13. Zhu Y, Tchkonia T, Pirtskhalava T, Gower AC, Ding H, Giorgadze N, et al. The Achilles’ heel of senescent cells: from transcriptome to senolytic drugs. Aging Cell. 2015;14(4):644–58.

14. Farr JN, Xu M, Weivoda MM, Monroe DG, Fraser DG, Onken JL, et al. Targeting cellular senescence prevents age-related bone loss in mice. Nat Med. 2017;23(9):1072–9.

15. Baker DJ, Wijshake T, Tchkonia T, LeBrasseur NK, Childs BG, van de Sluis B, et al. Clearance of p16Ink4a-positive senescent cells delays ageing-associated disorders. Nature. 2011;479(7372):232-6.

16. Baker DJ, Childs BG, Durik M, Wijers ME, Sieben CJ, Zhong J, et al. Naturally occurring p16(Ink4a)-positive cells shorten healthy lifespan. Nature. 2016;530(7589):184-9.

17. Hansen DV, Hanson JE, and Sheng M. Microglia in Alzheimer’s disease. J Cell Biol. 2018;217(2):459–72.

18. Nott A, Holtman IR, Coufal NG, Schlachetzki JCM, Yu M, Hu R, et al. Brain cell type-specific enhancer-promoter interactome maps and disease-risk association. Science. 2019;366(6469):1134-9.

19. Schwabe T, Srinivasan K, and Rhinn H. Shifting paradigms: The central role of microglia in Alzheimer’s disease. Neurobiol Dis. 2020;143:104962.

20. Kinney JW, Bemiller SM, Murtishaw AS, Leisgang AM, Salazar AM, and Lamb BT. Inflammation as a central mechanism in Alzheimer’s disease. Alzheimers Dement (N Y). 2018;4:575–90.

21. Asai H, Ikezu S, Tsunoda S, Medalla M, Luebke J, Haydar T, et al. Depletion of microglia and inhibition of exosome synthesis halt tau propagation. Nat Neurosci. 2015;18(11):1584–93.

22. Yoshiyama Y, Higuchi M, Zhang B, Huang SM, Iwata N, Saido TC, et al. Synapse loss and microglial activation precede tangles in a P301S tauopathy mouse model. Neuron. 2007;53(3):337–51.

23. Kitazawa M, Oddo S, Yamasaki TR, Green KN, and LaFerla FM. Lipopolysaccharide-induced inflammation exacerbates tau pathology by a cyclin-dependent kinase 5-mediated pathway in a transgenic model of Alzheimer’s disease. J Neurosci. 2005;25(39):8843–53.

24. Bowman GL, Dayon L, Kirkland R, Wojcik J, Peyratout G, Severin IC, et al. Blood-brain barrier breakdown, neuroinflammation, and cognitive decline in older adults. Alzheimers Dement. 2018;14(12):1640–50.

25. Nation DA, Sweeney MD, Montagne A, Sagare AP, D’Orazio LM, Pachicano M, et al. Blood-brain barrier breakdown is an early biomarker of human cognitive dysfunction. Nat Med. 2019;25(2):270–6.

26. Wei Z, Liu H, Lin Z, Yao M, Li R, Liu C, et al. Non-contrast assessment of blood-brain barrier permeability to water in mice: An arterial spin labeling study at cerebral veins. Neuroimage. 2023;268:119870.

27. Lin Z, Li Y, Su P, Mao D, Wei Z, Pillai JJ, et al. Non-contrast MR imaging of blood-brain barrier permeability to water. Magn Reson Med. 2018;80(4):1507–20.

28. Lin Z, Sur S, Liu P, Li Y, Jiang D, Hou X, et al. Blood-Brain Barrier Breakdown in Relationship to Alzheimer and Vascular Disease. Ann Neurol. 2021;90(2):227–38.

29. Costantini LC, Barr LJ, Vogel JL, and Henderson ST. Hypometabolism as a therapeutic target in Alzheimer’s disease. BMC Neurosci. 2008;9 Suppl 2(Suppl 2):S16.

30. Wei Z, Xu J, Liu P, Chen L, Li W, van Zijl P, et al. Quantitative assessment of cerebral venous blood T(2) in mouse at 11.7T: Implementation, optimization, and age effect. Magn Reson Med. 2018;80(2):521–8.

31. Riessland M, and Orr ME. Translating the Biology of Aging into New Therapeutics for Alzheimer’s Disease: Senolytics. J Prev Alzheimers Dis. 2023;10(4):633–46.

32. Nambiar A, Kellogg D, 3rd, Justice J, Goros M, Gelfond J, Pascual R, et al. Senolytics dasatinib and quercetin in idiopathic pulmonary fibrosis: results of a phase I, single-blind, single-center, randomized, placebo-controlled pilot trial on feasibility and tolerability. EBioMedicine. 2023;90:104481.

33. Bennett RE, Robbins AB, Hu M, Cao X, Betensky RA, Clark T, et al. Tau induces blood vessel abnormalities and angiogenesis-related gene expression in P301L transgenic mice and human Alzheimer’s disease. Proc Natl Acad Sci U S A. 2018;115(6):E1289–E98.

34. Tetrault S, Chever O, Sik A, and Amzica F. Opening of the blood-brain barrier during isoflurane anaesthesia. Eur J Neurosci. 2008;28(7):1330–41.

35. Abbott NJ, Patabendige AA, Dolman DE, Yusof SR, and Begley DJ. Structure and function of the blood-brain barrier. Neurobiol Dis. 2010;37(1):13–25.

36. Michalicova A, Majerova P, and Kovac A. Tau Protein and Its Role in Blood-Brain Barrier Dysfunction. Front Mol Neurosci. 2020;13:570045.

37. de Vries HE, Kooij G, Frenkel D, Georgopoulos S, Monsonego A, and Janigro D. Inflammatory events at blood-brain barrier in neuroinflammatory and neurodegenerative disorders: implications for clinical disease. Epilepsia. 2012;53 Suppl 6(Suppl 6):45–52.

38. Persidsky Y, Hill J, Zhang M, Dykstra H, Winfield M, Reichenbach NL, et al. Dysfunction of brain pericytes in chronic neuroinflammation. J Cereb Blood Flow Metab. 2016;36(4):794–807.

39. Zlokovic BV. The blood-brain barrier in health and chronic neurodegenerative disorders. Neuron. 2008;57(2):178–201.

40. Laurent C, Buee L, and Blum D. Tau and neuroinflammation: What impact for Alzheimer’s Disease and Tauopathies? Biomed J. 2018;41(1):21–33.

41. Bellucci A, Westwood AJ, Ingram E, Casamenti F, Goedert M, and Spillantini MG. Induction of inflammatory mediators and microglial activation in mice transgenic for mutant human P301S tau protein. Am J Pathol. 2004;165(5):1643–52.

42. Hickman SE, Kingery ND, Ohsumi TK, Borowsky ML, Wang LC, Means TK, et al. The microglial sensome revealed by direct RNA sequencing. Nat Neurosci. 2013;16(12):1896–905.

43. Keren-Shaul H, Spinrad A, Weiner A, Matcovitch-Natan O, Dvir-Szternfeld R, Ulland TK, et al. A Unique Microglia Type Associated with Restricting Development of Alzheimer’s Disease. Cell. 2017;169(7):1276–90 e17.

44. Krasemann S, Madore C, Cialic R, Baufeld C, Calcagno N, El Fatimy R, et al. The TREM2-APOE Pathway Drives the Transcriptional Phenotype of Dysfunctional Microglia in Neurodegenerative Diseases. Immunity. 2017;47(3):566–81 e9.

45. Brelstaff JH, Mason M, Katsinelos T, McEwan WA, Ghetti B, Tolkovsky AM, et al. Microglia become hypofunctional and release metalloproteases and tau seeds when phagocytosing live neurons with P301S tau aggregates. Sci Adv. 2021;7(43):eabg4980.

46. Chen Y, and Colonna M. Microglia in Alzheimer’s disease at single-cell level. Are there common patterns in humans and mice? J Exp Med. 2021;218(9).

47. Jiang D, Liu P, Li Y, Mao D, Xu C, and Lu H. Cross-vendor harmonization of T(2) - relaxation-under-spin-tagging (TRUST) MRI for the assessment of cerebral venous oxygenation. Magn Reson Med. 2018;80(3):1125–31.

48. Wei Z, Xu J, Chen L, Hirschler L, Barbier EL, Li T, et al. Brain metabolism in tau and amyloid mouse models of Alzheimer’s disease: An MRI study. NMR Biomed. 2021;34(9):e4568.

49. Wei Z, Chen L, Lin Z, Jiang D, Xu J, Liu P, et al. Optimization of phase-contrast MRI for the estimation of global cerebral blood flow of mice at 11.7T. Magn Reson Med. 2019;81(4):2566–75.

50. Aggarwal M, Mori S, Shimogori T, Blackshaw S, and Zhang J. Three-dimensional diffusion tensor microimaging for anatomical characterization of the mouse brain. Magn Reson Med. 2010;64(1):249–61.

51. Tournier JD, Smith R, Raffelt D, Tabbara R, Dhollander T, Pietsch M, et al. MRtrix3: A fast, flexible and open software framework for medical image processing and visualisation. NeuroImage. 2019;202:116137.

52. Aggarwal M, Zhang J, Miller MI, Sidman RL, and Mori S. Magnetic resonance imaging and micro-computed tomography combined atlas of developing and adult mouse brains for stereotaxic surgery. Neuroscience. 2009;162(4):1339–50.

