## supplemental Figure S1 to S5 for "Senolytic therapy preserves blood-brain barrier integrity and promotes microglia homeostasis in a tauopathy model"

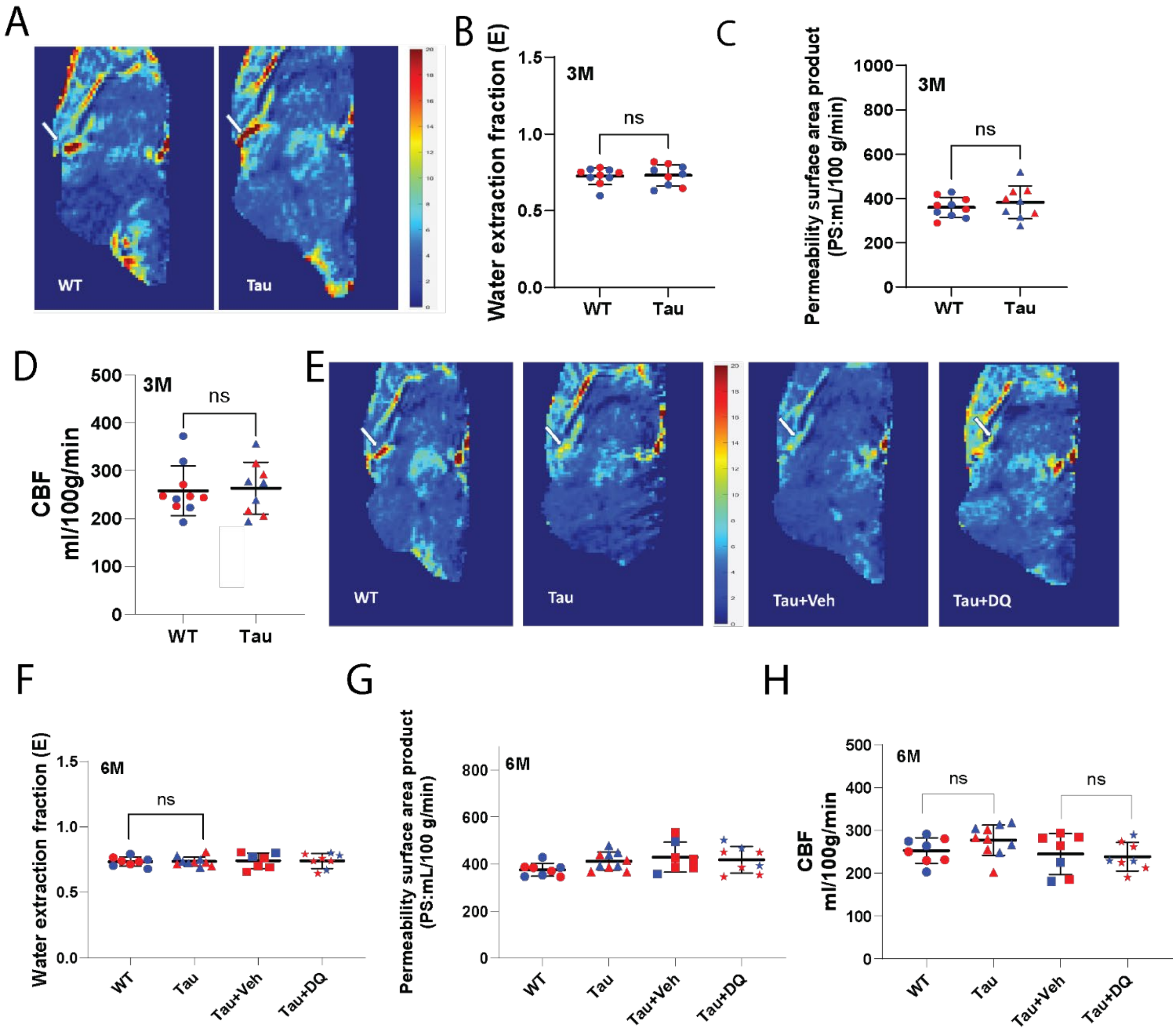


**Figure S1. Blood-brain barrier permeability measures in 3- and 6-month-old PS19 mice and their littermate wild type controls**. (**A**) Representative images of WEPCAST MRI images from 3 months old wild type (WT) control and PS19 (Tau) mice. (**B-C**) Water extraction fraction and permeability surface area product (PS) in 3 months old mice at indicated genotypes. n = 9-10 mice/group. ns = no statistical significance (**D**) Cerebral blood flow (CBF) measures in 3 months old mice. n = 9-10 mice/group. ns = no statistical significance. (**E**) Representative images of WEPCAST MRI images from 6 months old wild type (WT) control, PS19 (Tau) mice, PS19 mice treated with vehicle (Tau+Veh) or DQ (Tau+DQ). (**F-G**) Water extraction fraction and permeability surface area product (PS) in 6 months old mice at indicated groups. (**H**) Cerebral blood flow (CBF) measures in 6 months old mice. n = 7-10 mice/group; ns = no statistical significance by standard Student's test (B-D) or *ANOVA* with Bonferroni correction for multiple comparisons (F-H). In all graphs, individual data points are indicated; Red dots represent female mice and blue dots represent male mice. All data are shown as Mean and SD.

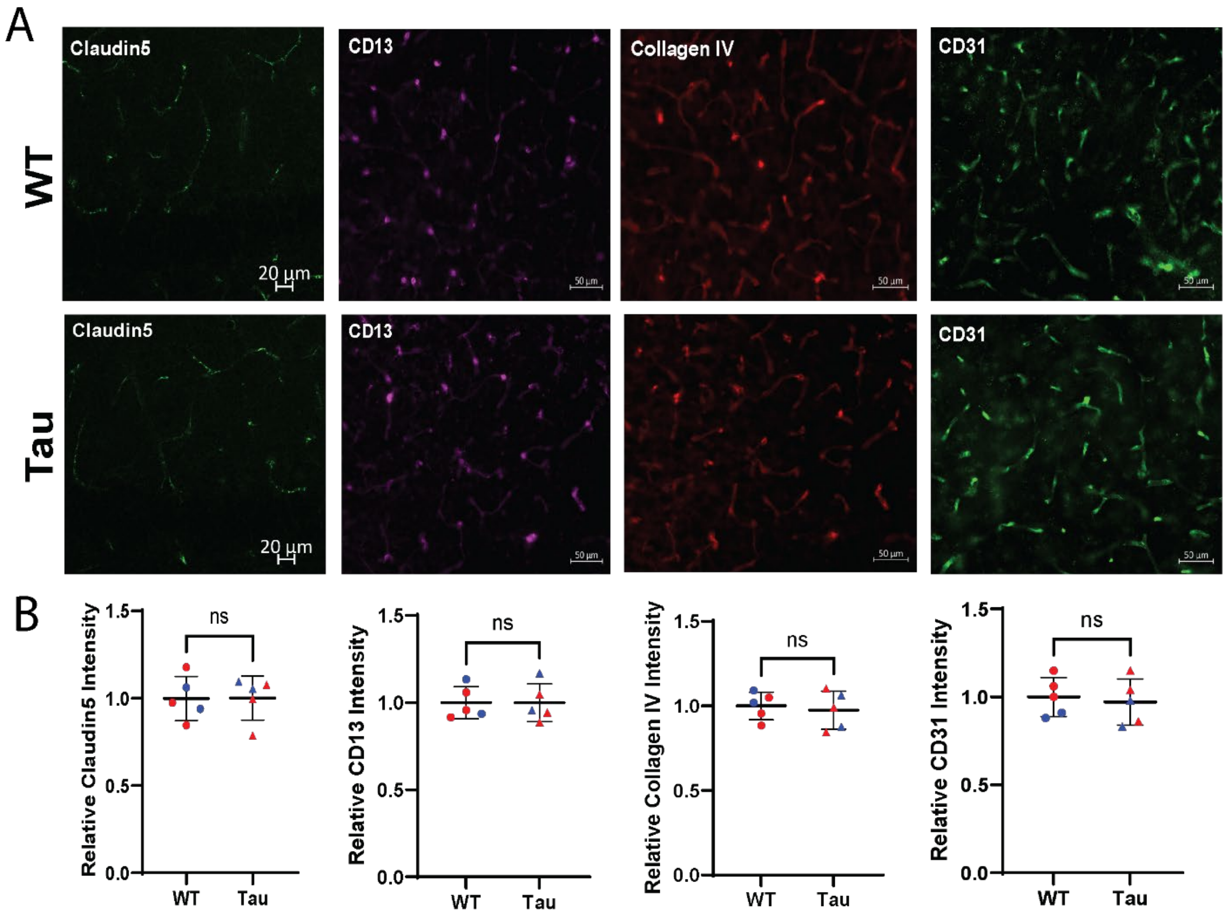


**Figure S2.** Immunohistochemistry of tight junction protein Claudin 5, pericyte marker protein CD13, vascular base membrane protein Collagen IV, and endothelia cell marker CD31 in 9-month-old wild type (WT) mice and PS19 mice (Tau). (A) representative images of immunostaining for indicated proteins. Scale bar for Claudin 5 is 20 μm; 50 μm for other three proteins. (B) Quantification of fluorescent intensity of indicated proteins. n=5. All data are shown as individual data points, Mean and SD. ns = no statistical significance between WT and PS19 (Tau) groups by Standard Student's *t*-test.

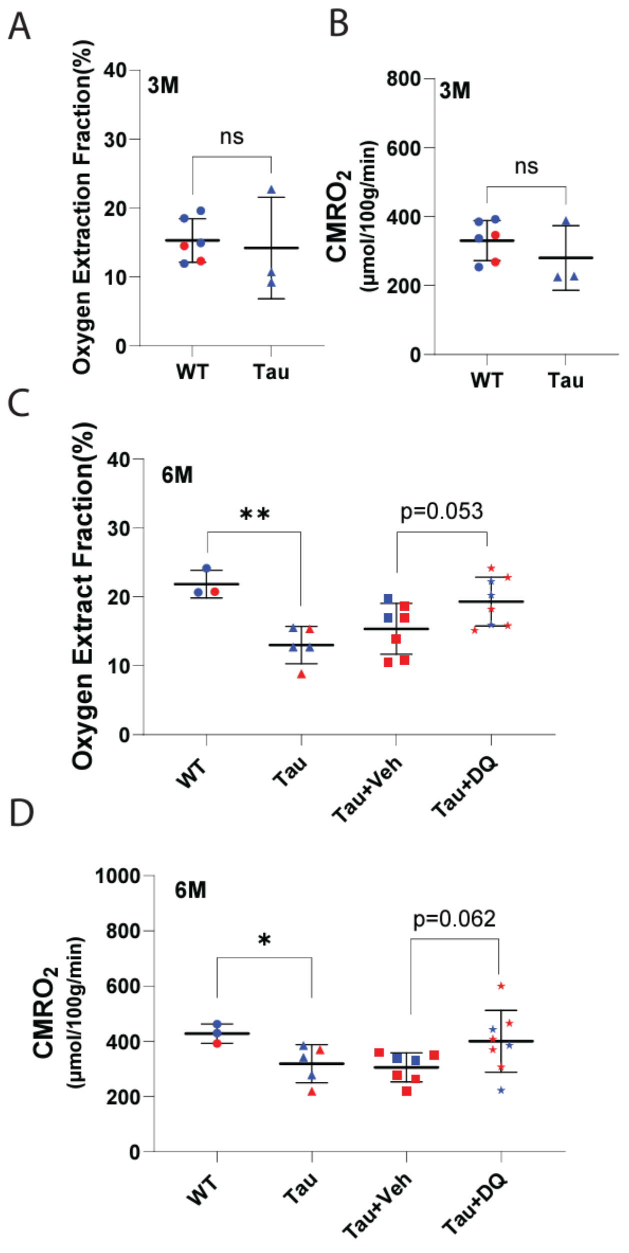


**Figure S3. Oxygen extraction fraction and global cerebral metabolic rate of oxygen (CMRO2) in 3 and 6 months old PS19 mice**. (**A-B**) Oxygen extraction fraction (OEF, A) and global cerebral metabolic rate of oxygen (CMRO2, B) were detected by noninvasive T2 relaxation under spin tagging (TRUST) and phase contrast MRI in 3 months old PS19 (Tau) mice and age-matched wild type (WT) controls. 6 mice in WT group and 3 mice in PS19 group. (**C-D**) OEF and CMRO2 measures in 6 months old mice at indicated groups. ns, no statistical significance. n = 3-8 mice/group; *$p<0.05$, **$p<0.01$ by standard Student's test (A-B) or *ANOVA* with Bonferroni correction for multiple comparisons (C-D). In all graphs, each individual data points are indicated; Red dots represent female mice and blue dots represent male mice. All data are shown as individual data points, Mean and SD.

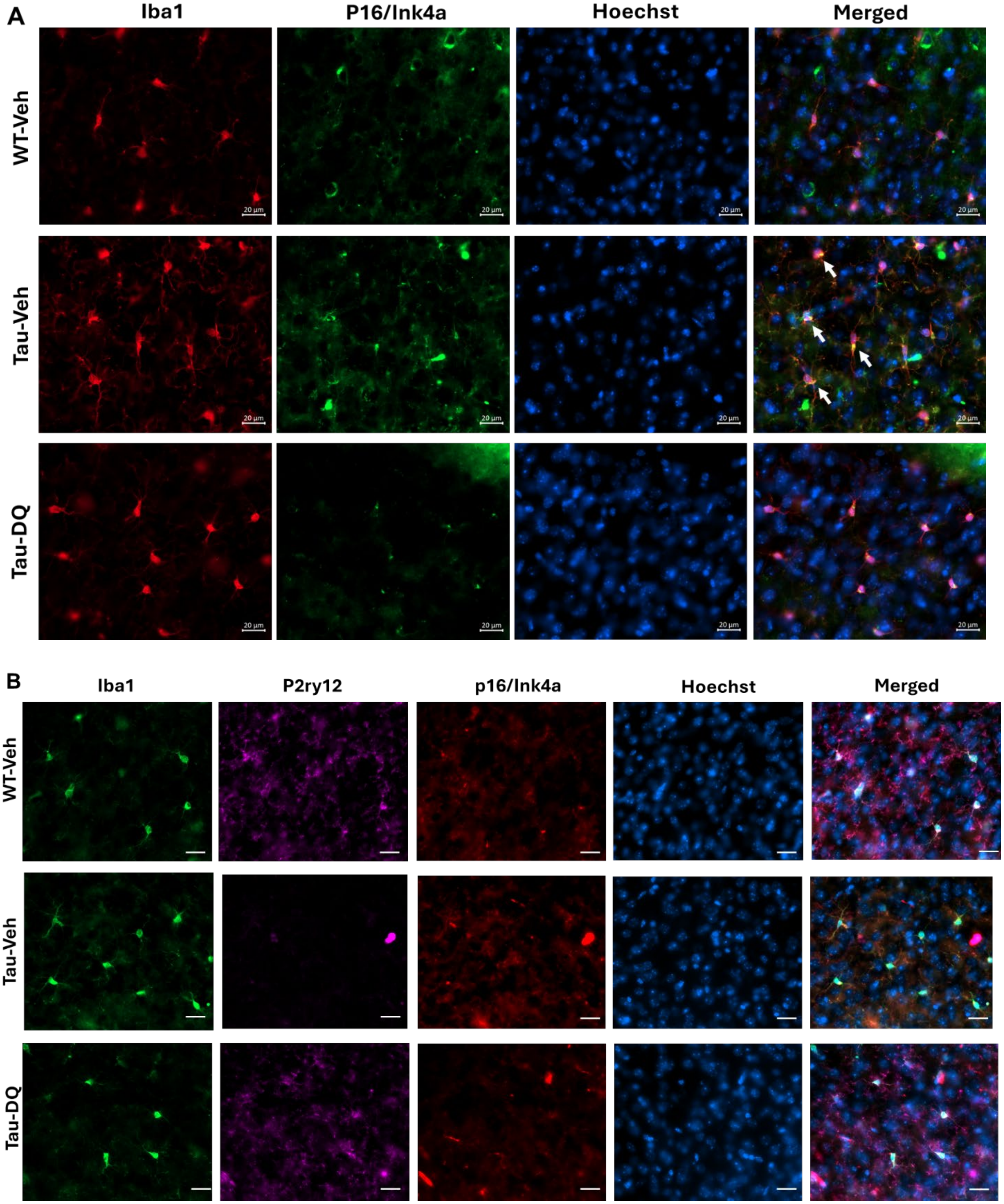
A
Iba1
P16/Ink4a
Hoechst
Merged
WT-Veh
Tau-Veh
Tau-DQ
20 µm
B
Iba1
P2ry12
p16/Ink4a
Hoechst
Merged
WT-Veh
Tau-Veh
Tau-DQ

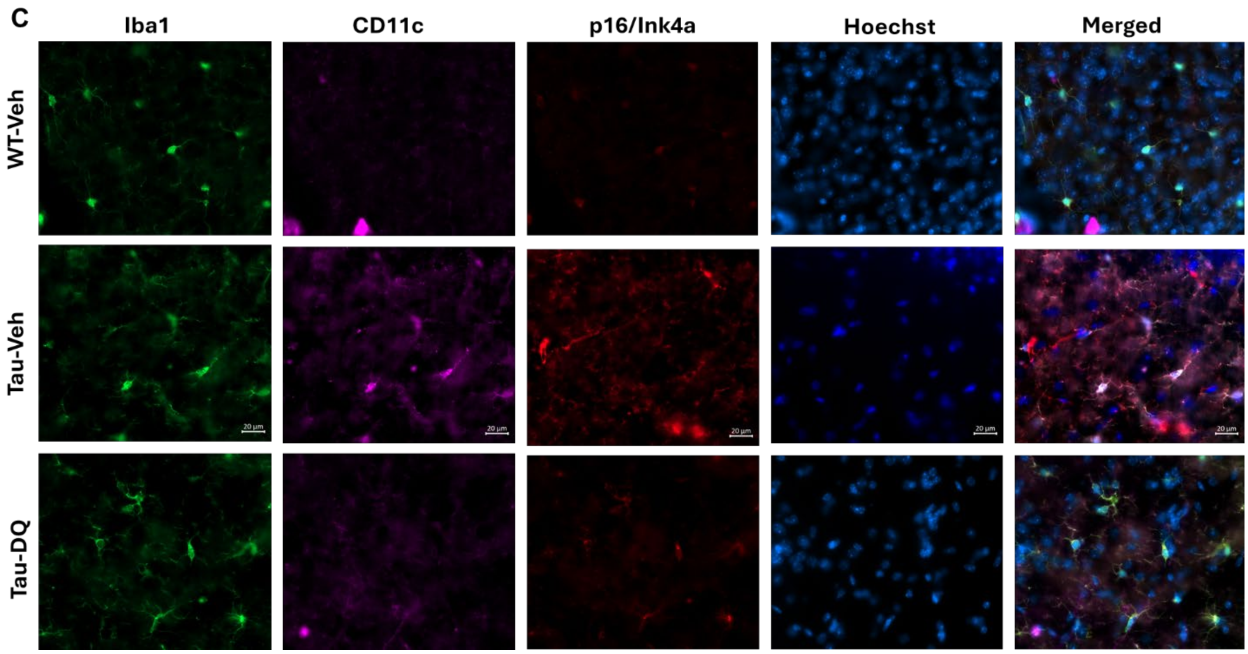


**Figure S4. Co-immunostaining of cell senescence marker p14/Ink4a with microglia marker protein Iba1 (A), p2ry12 (B), and CD11c (C) in the mouse brains from wild type mice with vehicle (WT-Veh), PS19 mice with vehicle (Tau-Veh), or PS19 mice with DQ treatment (Tau-DQ).** Note increased p16/Ink4a immunosignals in Tau-Veh mouse brain comparing to WT-Veh brain; and DQ treatment reduced p16/Ink4a immunosignals in PS19 mice.

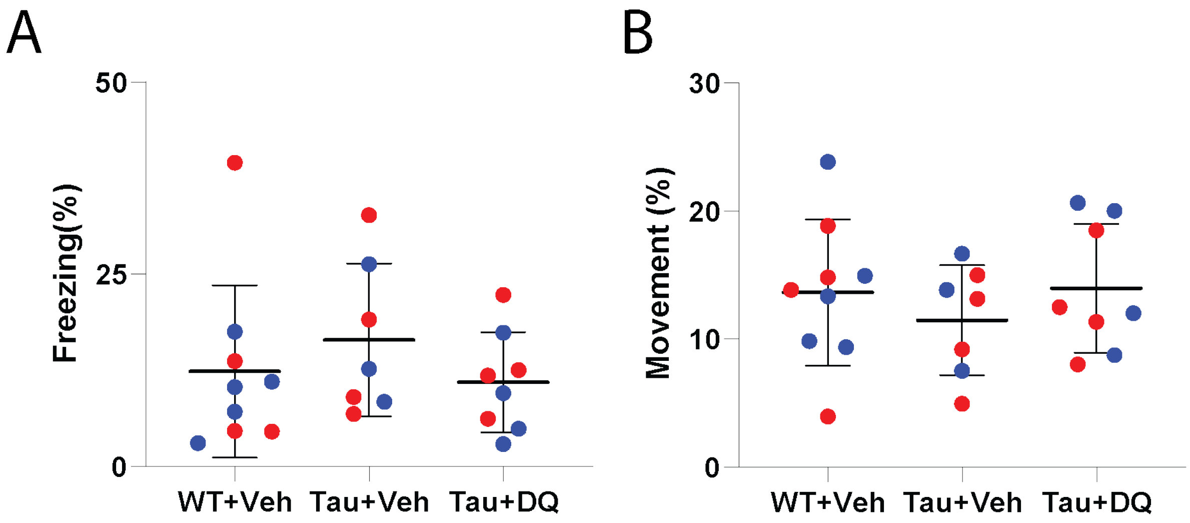


**Figure S5. Total movement and % freezing data from a non-stimulus time period among three groups.** There is no significant difference in total movement and % of freezing from a non-stimulus time period among three groups. *ANOVA* and student's t tests, $p > 0.05$ between any two groups.
